# The autophagic response to *Staphylococcus aureus* provides an intracellular niche in neutrophils

**DOI:** 10.1101/581223

**Authors:** Tomasz K. Prajsnar, Justyna J. Serba, Bernice M. Dekker, Josie F. Gibson, Samrah Masud, Angeleen Fleming, Simon A. Johnston, Stephen A. Renshaw, Annemarie H. Meijer

## Abstract

*Staphylococcus aureus* is a major human pathogen causing multiple pathologies, from cutaneous lesions to life-threatening sepsis. Although neutrophils contribute to immunity against *S. aureus*, multiple lines of evidence suggest that these phagocytes can provide an intracellular niche for staphylococcal dissemination. However, the mechanism of neutrophil subversion by intracellular *S. aureus* remains unknown. Targeting of intracellular pathogens by autophagy is recognised as an important component of host innate immunity, but whether autophagy is beneficial or detrimental to *S. aureus*-infected hosts remains controversial. Here, using larval zebrafish we show that *S. aureus* is rapidly decorated by the autophagy marker Lc3 following engulfment by macrophages and neutrophils. Upon phagocytosis by neutrophils, Lc3-positive, non-acidified spacious phagosomes are formed. This response is dependent on phagocyte NADPH oxidase as both *cyba* knockdown and diphenyleneiodonium (DPI) treatment inhibited Lc3 decoration of phagosomes. Importantly, NADPH oxidase inhibition diverted neutrophil *S. aureus* processing into tight acidified vesicles, which resulted in increased host resistance to the infection. Some intracellular bacteria within neutrophils were also tagged by p62-GFP fusion protein and loss of p62 impaired host defence. Taken together, we have shown that intracellular handling of *S. aureus* by neutrophils is best explained by Lc3-associated phagocytosis (LAP), which appears to provide an intracellular niche for bacterial pathogenesis, while the selective autophagy receptor p62 is host-protective. The antagonistic roles of LAP and p62-mediated pathways in *S. aureus*-infected neutrophils may explain the conflicting reports relating to anti-staphylococcal autophagy and provide new insights for therapeutic strategies against antimicrobial resistant staphylococci.

## INTRODUCTION

*Staphylococcus aureus* is a highly successful human pathogen causing a wide range of diseases [1]. This microorganism is a leading cause of fatal bacteraemia, with mortality rates reaching 30% [2] and multidrug resistance a particular concern [3]. Current therapeutic strategies to treat antimicrobial-resistant staphylococcal infections are becoming limited, and despite multiple attempts, there is still no vaccine available [4].

Although traditionally considered as an extracellular pathogen, accumulating evidence suggests that *S. aureus* is not only capable of inducing phagocyte lysis but also able to survive within professional phagocytes such as macrophages [5] and neutrophils [6]. Neutrophils, although shown to play a role in immunity against *S. aureus* [7], can also provide an intracellular niche for staphylococcal dissemination or persistence [8–11]. However, little is known of how *S. aureus* are able to subvert host cells to avoid phagocyte killing. A better understanding of the interactions of intracellular *S. aureus* with phagocytes is needed in order to develop therapies based on immunomodulatory approaches [12].

Autophagy is the evolutionarily-conserved intracellular degradation pathway by which eukaryotic cells scavenge their own cytoplasmic contents through sequestration into a nascent phagophore whose edges subsequently fuse to form a double membrane-surrounded vesicle called the autophagosome and which then fuses with the lysosome for degradation [13]. Microtubule-associated protein 1 light chain 3 (MAP1LC3, LC3) is an autophagosomal marker decorating membrane phagophores during elongation and in the resulting autophagosomes. In addition to nutrient-recycling functions, targeting of intracellular pathogens by the autophagic machinery has become recognised as an important component of host innate immunity in a process called xenophagy. This selective degradation requires the use of ubiquitin receptors such as p62, also named sequestosome 1 (SQSTM1). However, multiple intracellular pathogens such as *Mycobacterium tuberculosis, Salmonella* Typhimurium or *Listeria monocytogenes* have evolved strategies to inhibit or subvert the autophagic response [14].

LC3-associated phagocytosis (LAP) is a recently described process that is similar to, but functionally and molecularly distinct from, canonical autophagy and which lacks formation of the characteristic double-membrane autophagosome [15,16]. In this pathway, bacteria-containing single-membraned phagosomes are decorated with a lipidated form of LC3 which is directly coupled to the phagosomal membrane. This process requires the core autophagy machinery responsible for LC3 conjugation to lipids such as Autophagy-related Gene 5 (ATG5), but does not require early events of autophagosome initiation such as the unc-51 like autophagy activating kinase 1 (ULK1) [17,18]. In addition, LAP also requires NADPH oxidase activity and phagosomal ROS formation [18,19]. Although it is believed that LC3 on a phagosomal membrane facilitates subsequent fusion with lysosomes and degradation of pathogens, depending on the cellular background, LAP can either accelerate or delay phagosome fusion with lysosomes [20].

To date, studies of autophagy on non-professional phagocytes infected with *S. aureus* provide conflicting results, with the core autophagic machinery shown to be either detrimental [21] or beneficial [22] to the infected host cells. Schnaith *et al.* have shown that *S. aureus* is taken up within RAB7-positive phagosomes of mouse embryonic fibroblasts and subsequently is trapped within LC3-positive vesicles which serve as a niche for bacterial replication. That process is dependent on bacterial virulence factors regulated by staphylococcal Accessory Gene Regulator (*agr*). In these studies, inhibition of the core autophagy machinery by *atg5* knock-out was beneficial to the host cell as it led to reduction of intracellular staphylococci [21]. It has also been shown that the staphylococcal toxin α-haemolysin, which is positively regulated by *agr*, participates in the activation of the autophagic pathway within non-professional phagocytes [23,24]. In contrast, Neumann *et al.* more recently demonstrated the protective role of xenophagy in *S. aureus* infection of murine fibroblasts NIH/3T3, where intracellular bacteria are ubiquitinated leading to recruitment of selective autophagy receptors such as NDP52 or p62 and *atg5* knock-out leads to increased numbers of intracellular staphylococci [22]. In addition, the role of autophagic response to *S. aureus*, as beneficial or detrimental to the host, might be cell-type specific. Importantly, the autophagic response to *S. aureus* within macrophages and neutrophils has not been studied in detail, and it is currently unknown how the different processes that rely on autophagy components, such as xenophagy and LAP, are involved in the interaction of professional phagocytes with staphylococci during systemic infection.

In this study, using a well-established zebrafish model of staphylococcal systemic infection [10,25] accompanied by *in vivo* imaging of transgenic zebrafish, we explore the autophagic response to *S. aureus* within professional phagocytes; macrophages and neutrophils. Our results identify LAP as the important pathway responsible for staphylococcal subversion of infected neutrophils leading to progression of systemic disease.

## RESULTS

### *S. aureus* is contained within Lc3-positive vesicles in both zebrafish macrophages and neutrophils

Due to its central role in autophagy and related responses, a lipidated form of the LC3 protein has been widely used as a marker for activation of the autophagy machinery in both mammalian and zebrafish systems [26,27]. Therefore, in order to study the autophagic response to *S. aureus* infection at the cellular level, we used a transgenic zebrafish line *Tg(CMV:EGFP-map1lc3b)zf155* [27], hereafter called CMV:GFP-Lc3. Embryos of this line were infected with mCherry-labelled *S. aureus* SH1000 [28] as previously described [25]. Infected CMV:GFP-Lc3 zebrafish were fixed at 1, 2 and 4 hours post infection (hpi) and confocal microscopy performed to visualise the formation of Lc3-positive vesicles associated with bacteria within infected phagocytes. GFP-Lc3 associations with bacteria within phagocytes were observed (Fig. 1a) with most Lc3-bacteria associations within infected phagocytes seen at 1 hour post infection (hpi) and reducing by 4 hpi (Fig. 1b).

**Figure 1:**
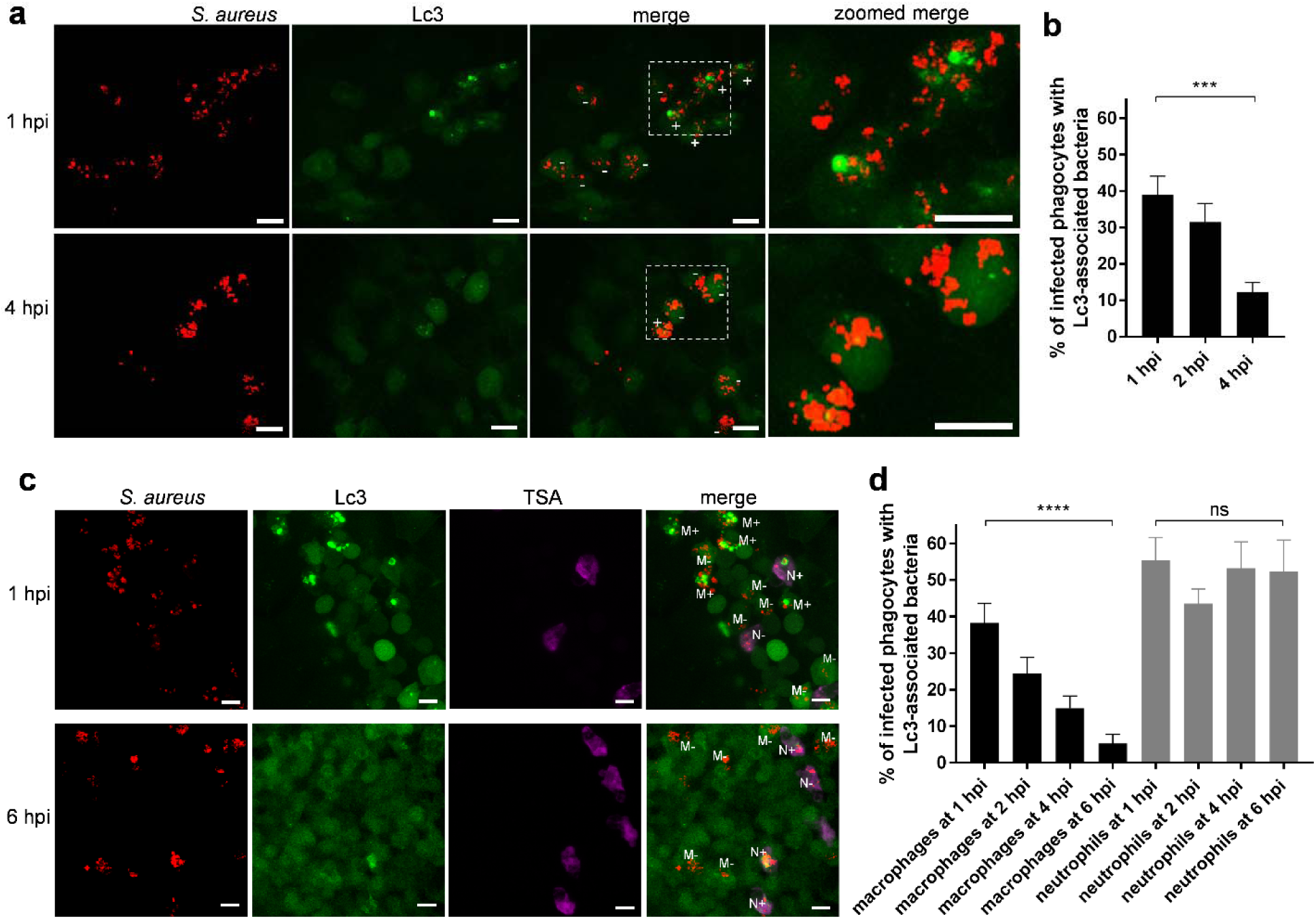
Different kinetics of the Lc3-mediated response in macrophages and neutrophils infected by *S. aureus*. a. Confocal photomicrographs shown as maximum intensity projections of CMV:GFP-Lc3 embryos infected with mCherry-labelled *S. aureus* at 1 (top panel) and 4 hpi (bottom panel). Phagocytes are seen containing bacteria with (+) or without (-) Lc3 aggregates. Images shown are representative of three independent experiments. Scale bars represent 10 µm.
b. Quantification of Lc3 associations with intracellular *S. aureus* at 1, 2 and 4 hpi in CMV:GFP-Lc3 embryos. Data are shown as mean +/− standard error of the mean (SEM) obtained from three independent experiments. *** *P*<0.001.
c. Confocal photomicrographs shown as maximum intensity projections of CMV:GFP-Lc3 transgenic embryos infected with mCherry-labelled *S. aureus.* Embryos were fixed at 1 (top panel) and 6 hpi (bottom panel) and chemically stained for Mpx activity (TSA, magenta). TSA-negative macrophages are seen containing bacteria with (M+) or without (M-) Lc3 aggregates as well as TSA-positive neutrophils containing bacteria with (N+) and without Lc3 aggregates (N-). The images shown are representative of three independent experiments. Scale bars represent 10 µm.
d. Quantification of Lc3 associations with intracellular *S. aureus* at 1, 2, 4 and 6 hpi within macrophages (black bars) and neutrophils (grey bars). Data are shown as mean +/− standard error of the mean (SEM) obtained from three independent experiments. **** *P*<0.0001, ns – not significant.

Subsequently, in order to characterise the observed Lc3-mediated response specifically in infected macrophages and neutrophils, fixed CMV:GFP-Lc3 embryos were subjected to histochemical staining for endogenous peroxidase activity [25] to distinguish neutrophils from macrophages, which are peroxidase negative in zebrafish [29]. In agreement with previous work, both neutrophils and macrophages were observed to take up *S. aureus*, with most bacteria detected within TSA-negative infected macrophages [25]. The number of Lc3-*S.aureus*-positive macrophages were seen to decrease over time (Fig. 1c, d), while the Lc3 associations with *S. aureus* within TSA-positive neutrophils remained high for up to 6 hpi. The Lc3 signal in infected neutrophils typically appeared in circular patterns around bacterial clusters, suggesting that it labels the vesicles containing *S. aureus*. This demonstrates that, although both macrophages and neutrophils are able to mount an Lc3-mediated response to *S. aureus* infection, the kinetics of this response differs.

Infected neutrophils represent a minority of infected phagocytes in systemically-infected embryonic zebrafish. Therefore, to study the neutrophil response specifically, we knocked down *irf8*, which leads to preferential development of neutrophils at the expense of macrophages [30]. This is a useful approach to manipulate neutrophil:macrophage ratios and has been successfully used in several infection studies [10,31–33]. This approach revealed that, as in embryos with normal myeloid cell ratios, around 55% of infected neutrophils still contained Lc3-associations with bacteria at 6 hpi (Fig. S1). Taken together, these data suggest that processing of Lc3-positive vesicles [18] is delayed in neutrophils compared to macrophages, suggesting a potential inhibition of autophagic flux within *S. aureus*-infected neutrophils.

### A neutrophil-specific autophagy reporter line confirms the Lc3-mediated response to *S. aureus* in neutrophils

We speculated that macrophages are able to use the machinery of autophagy to aid in intracellular processing of *S. aureus* bacteria, while neutrophils are less able to do so, potentially due to manipulation of autophagy by intracellular staphylococci. In order to study the neutrophil-specific Lc3-mediated response without the need for additional staining, we generated a transgenic zebrafish line where zebrafish Lc3 is fused with a tandem fluorophore RFP-GFP [34] and is expressed under a neutrophil-specific promoter *lyz* [35]: *Tg(lyz:RFP-GFP-map1lc3b)sh383* transgenic line (Fig. 2a), hereafter called *lyz*:RFP-GFP-Lc3.

**Figure 2:**
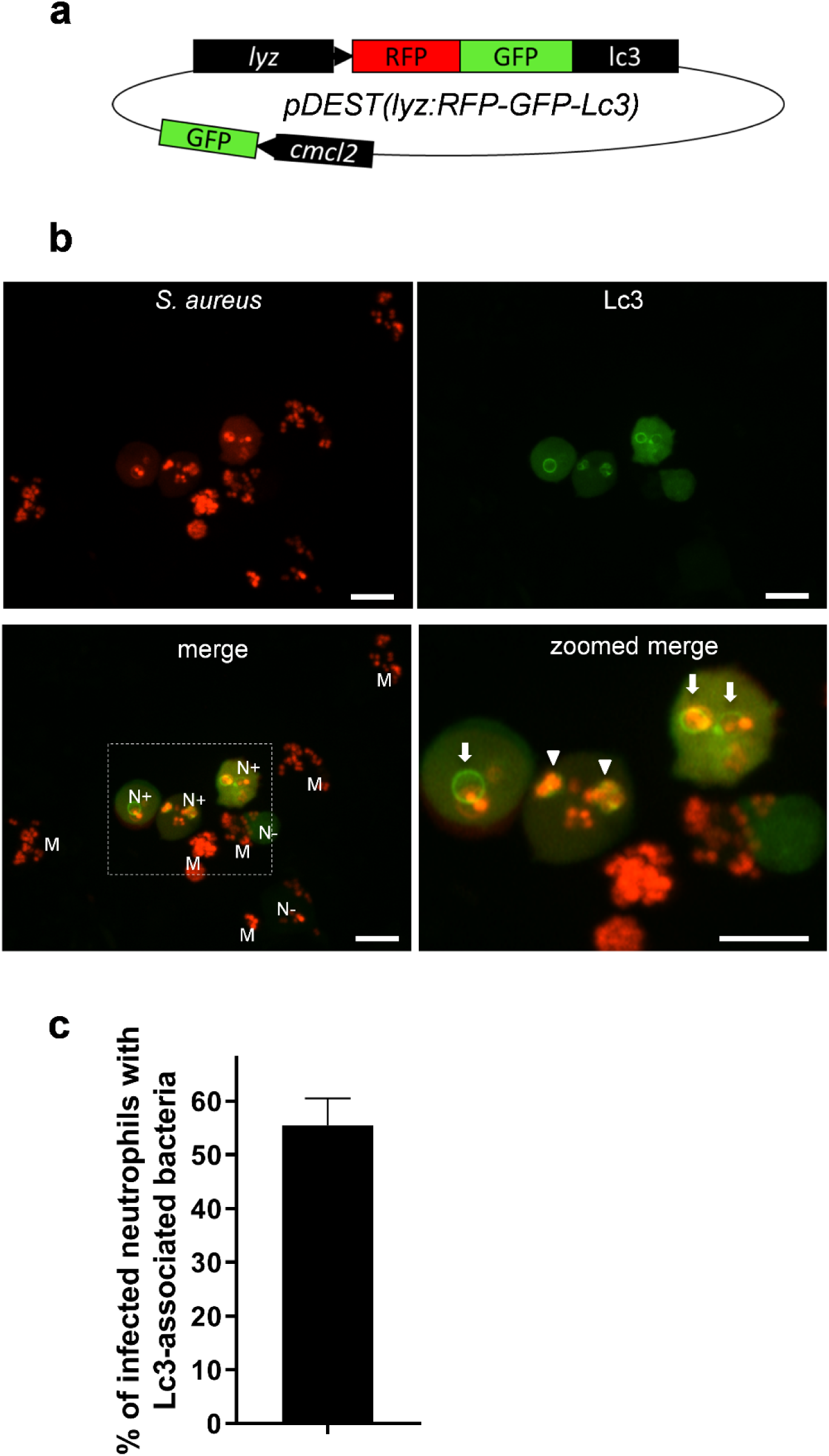
Generation of a *lyz*:RFP-GFP-Lc3 transgenic line in zebrafish confirms the Lc3-mediated response to *S. aureus* within neutrophils. a. Schematic of the pDEST(*lyz*:RFP-GFP-Lc3) construct encoding the fusion RFP-GFP-Lc3 protein under the neutrophil-specific *lyz* promoter. In addition, the heart marker *cmlc2*-driven GFP is used to facilitate the screening of the positive larvae.
b. Confocal images at maximum projection of the Lc3-mediated response at 1 hpi in live *lyz*:RFP-GFP-Lc3 embryos infected with mCherry-labelled *S. aureus*. Lyz-positive neutrophils are seen containing bacteria with (N+) or without (N-) Lc3 aggregates. Lyz-negative macrophages are also seen containing bacteria (M). The images shown are representative of three independent experiments. Arrows indicate spacious Lc3-positive vesicles, whereas arrowheads show tightly wrapped Lc3-associated bacteria. Scale bars represent 10 µm.
c. Quantification of Lc3 associations with intracellular *S. aureus* at 1 hpi within infected neutrophils of *lyz*:RFP-GFP-Lc3 embryos. Data are shown as mean +/− standard error of the mean (SEM) obtained from three independent experiments.

Upon systemic infection of *lyz*:RFP-GFP-Lc3 transgenic embryos with mCherry-labelled *S. aureus*, Lc3-*S.aureus* associations at 1 hpi were identified in approximately 55% of infected *lyz*-positive neutrophils (Fig. 2b, c). In addition, as expected, a majority of internalised bacteria were found in *lyz*-negative (hence unlabelled) macrophages (Fig 2b). These results demonstrate that the *lyz*:RFP-GFP-Lc3 line provides a new tool for studying the Lc3-mediated response within neutrophils and allows high quality live imaging without the visible fluorescence of other cells as observed in the CMV:GFP-Lc3 line (Fig. 1). Furthermore, we could observe the fluorescent signal encircling spacious bacteria-containing vesicles, confirming that neutrophils mount an Lc3-mediated response to *S. aureus* infection (Fig. 2b).

### Functional NADPH oxidase is required for the formation of Lc3 aggregates associated with phagocytosed staphylococci

We have shown that that the transgenic Lc3 markers used in this study label bacteria-containing vesicles, suggesting these are formed by either selective autophagy or Lc3-associated phagocytosis (LAP). To distinguish these possibilities, we manipulated phagosomal ROS production, which has been shown to be specifically required for LAP [18,19,33,36]. Using well-validated morpholino-modified antisense oligonucleotide injection [37], we knocked down expression of *cyba* (a membrane bound subunit of phagocyte NADPH oxidase). Loss of Cyba led to a near complete absence of Lc3-*S.aureus* association in both macrophages and neutrophils of CMV:GFP-Lc3 embryos (Figure 3a, b). In agreement, infection of *cyba* knockdown *lyz*:RFP-GFP-Lc3 embryos led to no Lc3-*S.aureus* associations (Fig. 3c, d). In addition, Lc3-bacteria associations were lost upon genetic (Fig. 4a-c) and diphenyleneiodonium (DPI)-mediated chemical inhibition of NADPH oxidase (Fig. 4d-f) in macrophage-depleted, neutrophil-enriched (*irf8* knockdown) larvae. Therefore, these results suggest that Cyba and hence the NADPH oxidase complex plays an important role in the recruitment of Lc3 to *S. aureus* in both macrophages and neutrophils and therefore we propose that this response represents LAP.

**Figure 3:**
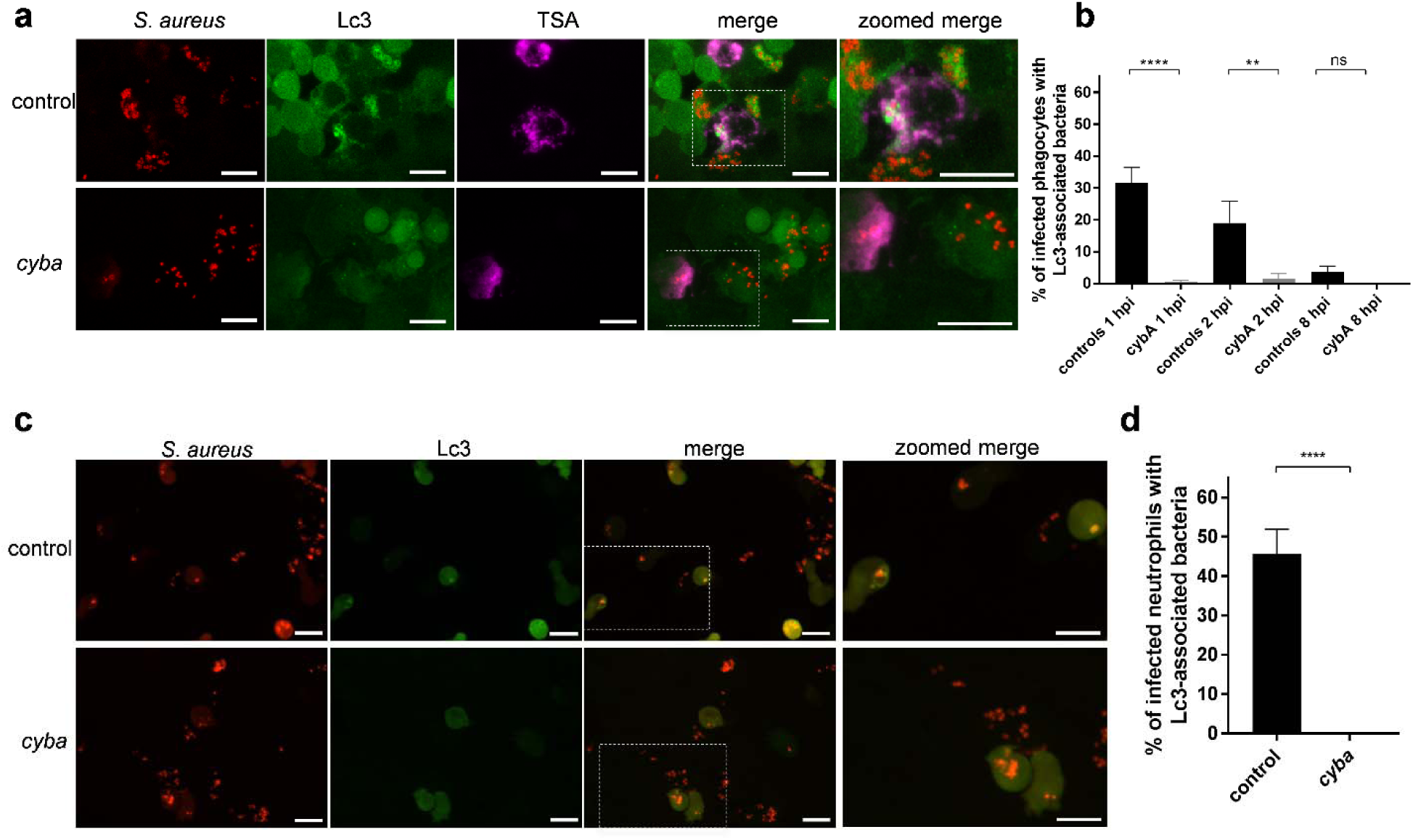
NADPH oxidase is required for the Lc3-mediated response to *S. aureus* infection. a. Confocal photomicrographs shown as maximum intensity projections of the Lc3-mediated response in control (top panel) and *cyba* knockdown (bottom panel) of CMV:GFP-Lc3 embryos infected with mCherry-labelled *S. aureus*. Embryos were fixed at 1 hpi and chemically stained for Mpx activity (TSA, magenta). Scale bars represent 10 µm.
b. Quantification of Lc3 associations with intracellular *S. aureus* at 1, 2 and 8 hpi within infected phagocytes of CMV:GFP-Lc3 control and *cyba* knockdown embryos. Data are shown as mean +/− standard error of the mean (SEM) obtained from three independent experiments. **** *P*<0.0001, ** *P*<0.01, ns – not significant.
c. Confocal photomicrographs shown as maximum intensity projections of the Lc3-mediated response at 1 hpi in control (top panel) and *cyba* knockdown live *lyz*:RFP-GFP-Lc3 embryos infected with mCherry-labelled *S. aureus*. Scale bars represent 10 µm.
d. Quantification of Lc3 associations with intracellular *S. aureus* at 1 hpi within infected neutrophils of Tg(lyz:GFP.RFP.Lc3) embryos. Data are shown as mean +/− standard error of the mean (SEM) obtained from three independent experiments. **** *P*<0.0001).

**Figure 4:**
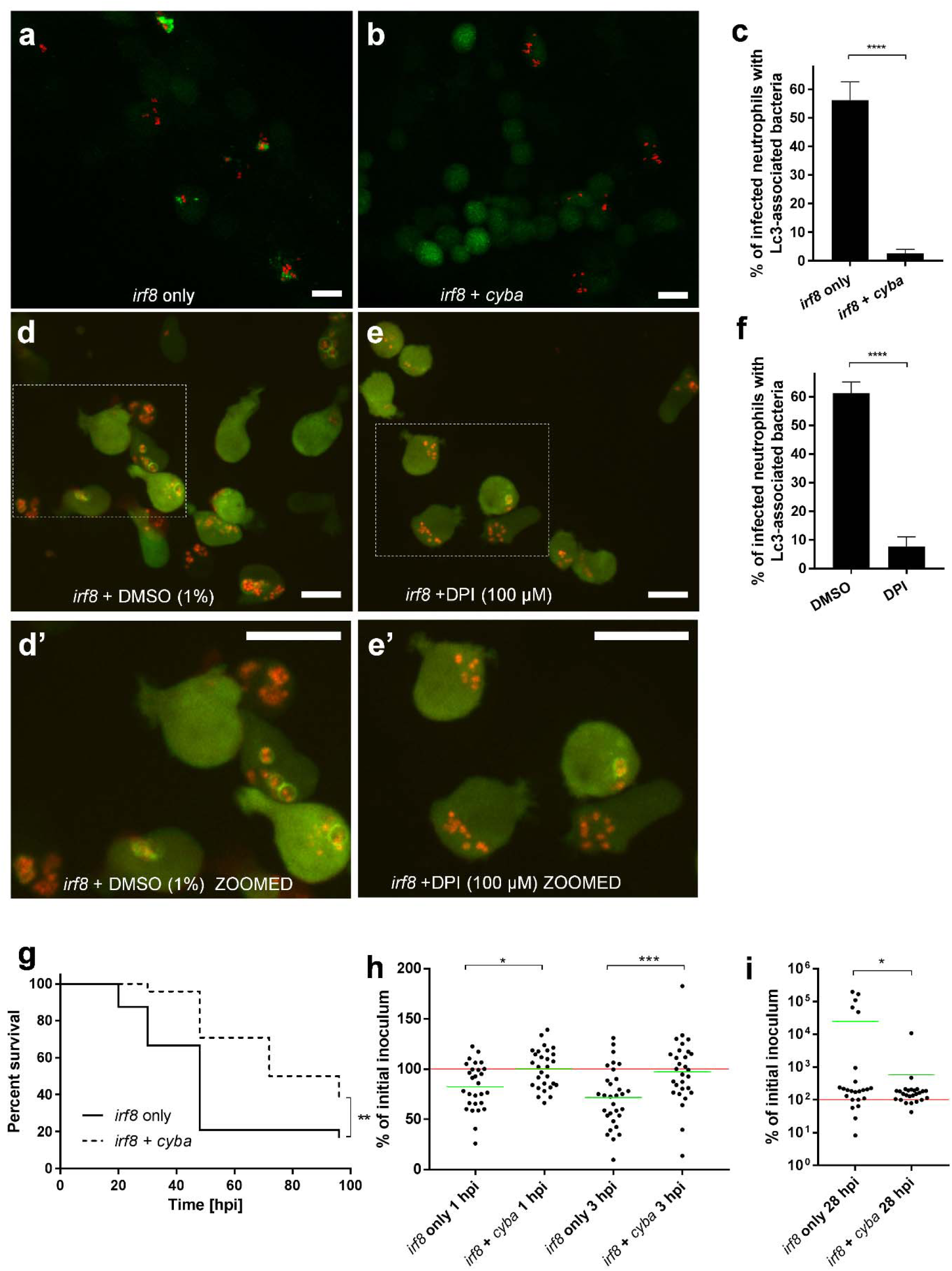
Formation of NADPH oxidase-dependent Lc3-positive vesicles containing *S. aureus* in neutrophils is detrimental for the infected host. a, b. Confocal photomicrographs shown as maximum intensity projections of the Lc3-mediated response at 1 hpi in *irf8* only (a) and *irf8* + *cyba* knockdown (b) CMV:GFP-Lc3 embryos infected with mCherry-labelled *S. aureus*. Scale bars represent 10 µm.
c. Quantification of Lc3 associations with intracellular *S. aureus* at 1 hpi within infected neutrophils of *irf8* only and *irf8* + *cyba* knockdown CMV:GFP-Lc3 embryos. **** *P*<0.0001.
d, e. Confocal photomicrographs shown as maximum intensity projections of the Lc3-mediated response at 1 hpi in control (DMSO) (d) and DPI-treated (e) *irf8* knockdown *lyz*:RFP-GFP-Lc3 embryos infected with mCherry-labelled *S. aureus*. Scale bars represent 10 µm.
d’, e’. Zoomed-in fragments of photomicrographs d and e.
f. Quantification of Lc3 associations with intracellular *S. aureus* at 1 hpi within infected neutrophils of control (DMSO) and DPI-treated *irf8* knockdown *lyz*:RFP-GFP-Lc3 embryos. **** *P*<0.0001.
g. Survival of *irf8* only or *irf8* + *cyba* knockdown zebrafish larvae following intravenous injection with *S. aureus* at 30 hpf (n≥25). This result is representative of three independent experiments. ** *P*<0.01.
h and i. The CFU counts of the *irf8* only or *irf8* + *cyba* knockdown larvae infected intravenously with *S. aureus* at 1 and 3 hpi (h) or 28 hpi (i). At each timepoint larvae were sacrificed, homogenised and the recovered staphylococci were enumerated by serial dilutions. The red line represents the level of the initial inoculum whereas the green lines represent the mean value of each group. * *P*<0.05, *** *P*<0.001.

To further determine what effect the inhibition of NADPH oxidase activity and the associated Lc3 response have on the staphylococcal pathogenesis in the neutrophil-enriched (*irf8* knockdown) zebrafish, a survival experiment with *S. aureus* infected embryos was performed. Strikingly, the survival of infected zebrafish with genetically or pharmacologically inhibited NADPH oxidase was higher than controls (Fig. 4g and S4a), suggesting that formation of NADPH oxidase-mediated Lc3-positive vesicles containing *S. aureus* in neutrophils is detrimental for the infected host. The observed difference in host survival prompted us to enumerate bacteria within larvae following infection. We found that within the first 3 hours of infection the neutrophils of *cyba/irf8* double knockdown embryos are significantly less proficient in killing the internalised bacteria than *irf8* only knockdown embryos (Fig. 4h). However, in line with the survival curves (Fig 4g), at the later timepoint (28 hpi) higher numbers of staphylococci were found in a subset of *irf8* only knockdown embryos compared with the *cyba/irf8* double knockdown embryos (Fig. 4i). Therefore, the early reduction in killing of internalised *S. aureus* by *cyba* deficient embryos ultimately resulted in increased host survival and less *in vivo* bacteria in infected embryos at 28 hpi. To corroborate the results obtained with the *irf8* knockdown, we used a second strategy to explore the impact of macrophage ablation (clodronate-containing liposomes) to specially deplete macrophages whilst not affecting neutrophils [38]. Similar to *irf8* knockdown, the *cyba* deficient, macrophage-depleted larvae were more resistant to *S. aureus* than their respective controls (Fig. S4b). Interestingly, the loss of *cyba* had no effect on the survival of infected larvae in the presence of macrophages (Fig. S4c), suggesting that NADPH oxidase-mediated processing of *S. aureus* in zebrafish macrophages does not play a vital role in intracellular handling of bacteria. We conclude that formation of NADPH oxidase-mediated, Lc3-positive vesicles containing *S. aureus* in neutrophils is detrimental for the infected host.

### NADPH oxidase-dependent neutrophil response to *S. aureus* forms spacious non-acidified Lc3-positive phagosomes

Since the Lc3 was seen to persist on phagosomes in infected neutrophils, we decided to further characterise this potentially host-detrimental response and assess the pH status of internalised bacteria. In order to determine whether live bacteria are required to induce a Lc3-mediated response within neutrophils, we injected heat-killed staphylococci into *lyz*:RFP-GFP-Lc3 embryos. This resulted in similar levels of Lc3-bacteria associations within infected neutrophils when compared to neutrophils infected with live bacteria (Fig. 5a-c) suggesting that the observed Lc3-mediated response does not require bacterial-driven damage of the phagosome or subsequent bacterial escape into the cytoplasm, similar to what has been recently proposed for LC3 recruitment to *Listeria monocytogenes* [39]. However, only live bacteria led to formation of spacious Lc3-positive phagosomes within infected neutrophils by 1 hpi (Fig. 5d). Therefore, these characteristic spacious phagosomes in *S. aureus*-infected neutrophils may be indicative of bacterial pathogenesis.

**Figure 5:**
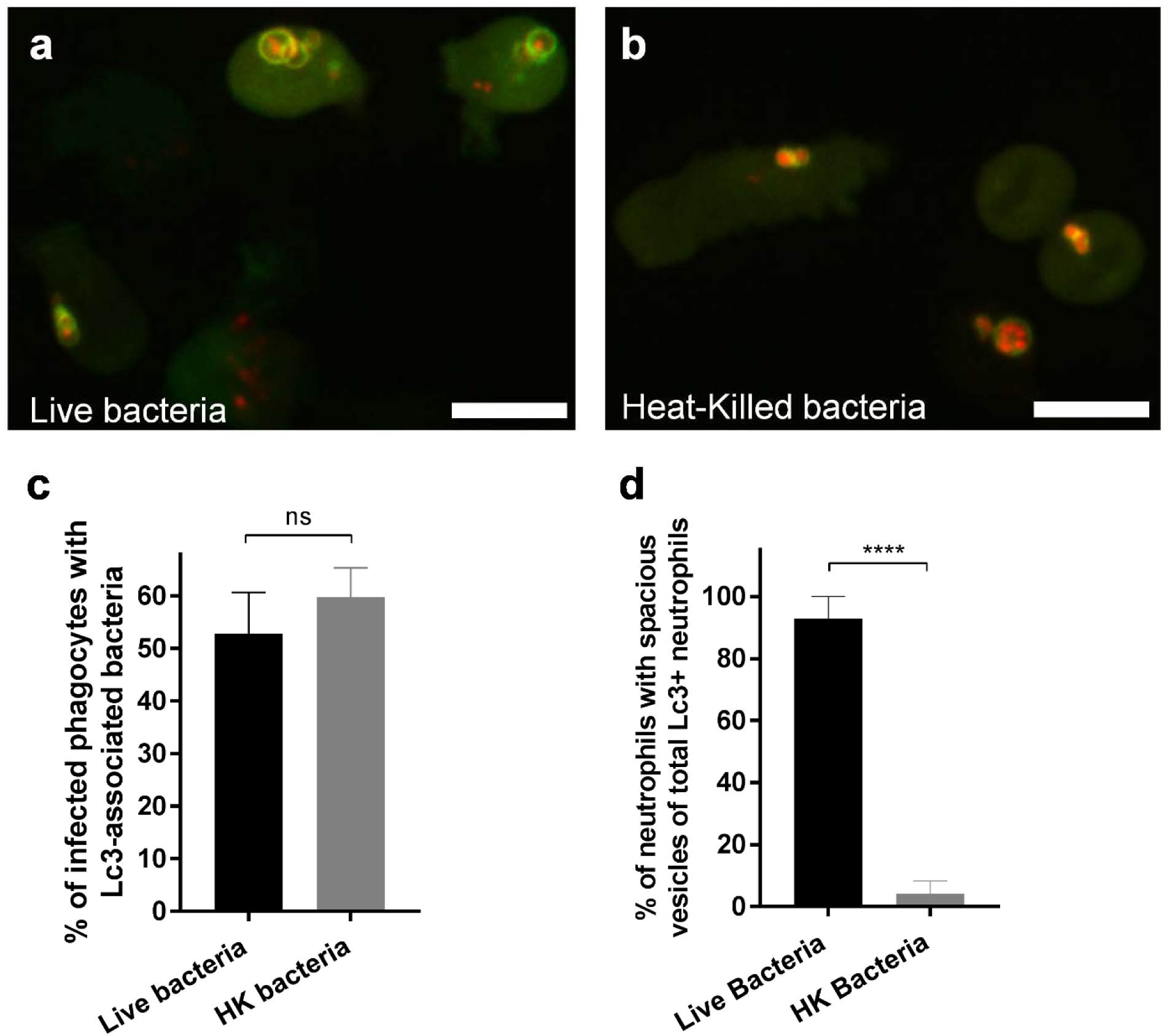
The Lc3-mediated response in neutrophils occurs to both live and heat-killed *S. aureus*, but spacious Lc3-positive vesicles are formed only with live bacteria. a, b. Confocal photomicrographs shown as maximum intensity projections of the Lc3-mediated response at 1 hpi in *lyz*:RFP-GFP-Lc3 embryos infected with live (a) or heat-killed (HK) (b) mCherry-labelled *S. aureus*. Scale bars represent 10 µm.
c. Quantification of Lc3 associations with intracellular *S. aureus* at 1 within neutrophils of *lyz*:RFP-GFP-Lc3 embryos infected with live or heat-killed mCherry-labelled *S. aureus*. ns – not significant.
d. Quantification of neutrophils with spacious *S. aureus*-containing phagosomes at 1 hpi within *lyz*:RFP-GFP-Lc3 embryos infected with live or heat-killed mCherry-labelled *S. aureus*. **** *P*<0.0001.

To assess whether ingested *S. aureus* were in acidic compartments, they were stained prior to inoculation with a combination of pH-sensitive dyes - pHrodo red and fluorescein succinimidyl esters (bacteria are green in neutral pH and red in acidic pH). While bacteria in control neutrophils were not in acidified compartments early during infection, the proportion of bacteria in acidified compartments increased slightly at later stages of infection (Fig. 6a, c). In *cyba* knockdown neutrophils, significantly more bacteria localised in acidic compartments (Fig. 6b, c). Importantly, the spacious Lc3-positive compartments containing bacteria in control fish remained at neutral pH (Fig. 6a). Therefore, lack of acidification of such phagosomes can potentially provide a non-acidified intraphagocyte niche for bacterial persistence or replication.

**Figure 6:**
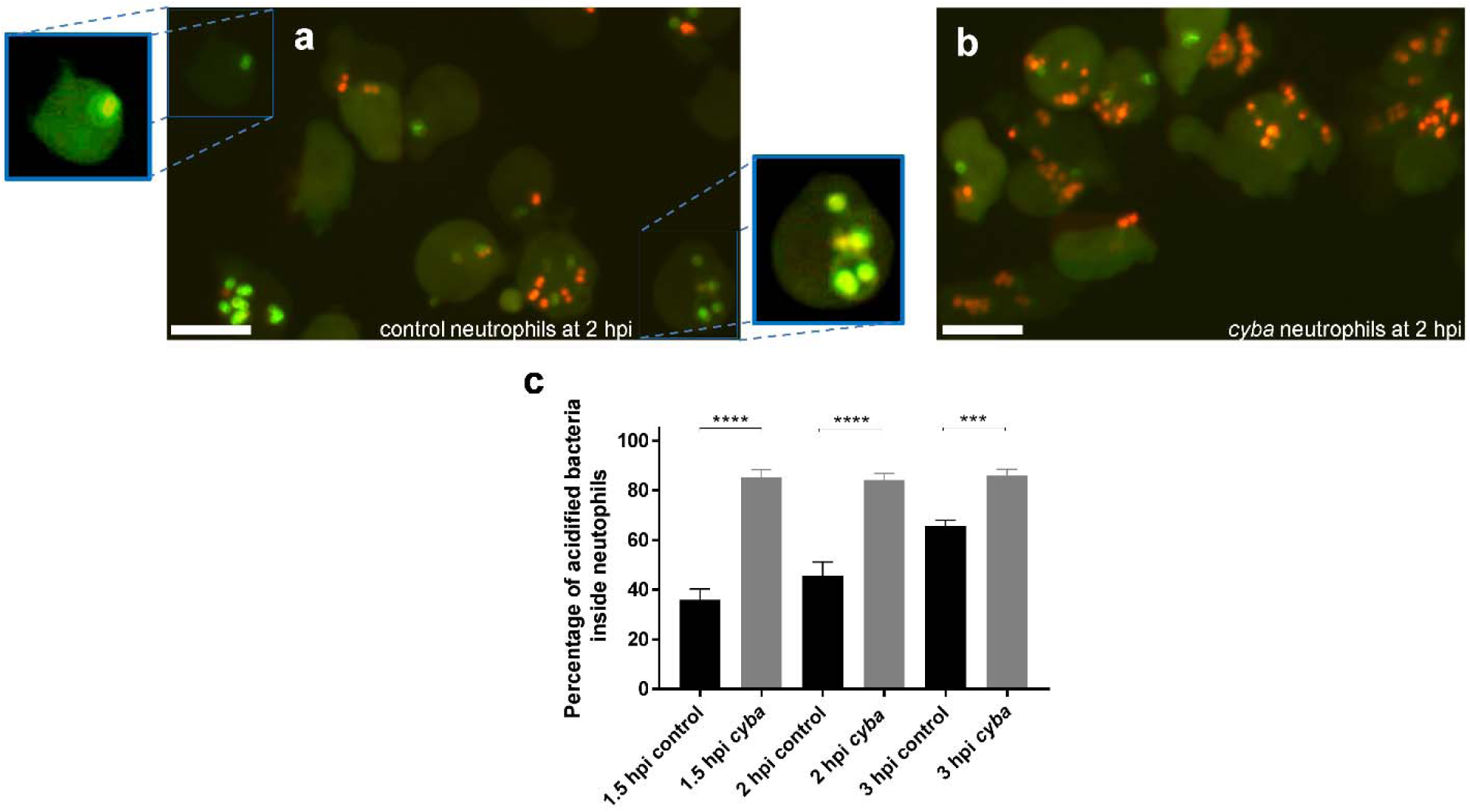
Loss of Cyba leads to increased acidification of neutrophil-ingested *S. aureus*. a, b. Confocal photomicrographs shown as maximum intensity projections of the control (left panel) and *cyba* knockdown (right panel) *lyz*:RFP-GFP-Lc3 embryos infected with *S. aureus* stained with pHrodo Red and Fluorescein pH-indicating dyes. Zoomed areas concentrate on neutrophils with spacious Lc3-positive phagosomes containing non-acidified staphylococci. Scale bars represent 10 µm.
c. Quantification of acidification rates at 1.5, 2 and 3 hpi of intracellular *S. aureus* within control and *cyba* knockdown neutrophils of embryos infected *S. aureus*. Data are shown as mean +/− standard error of the mean (SEM) obtained from three independent experiments. *** P<0.001, **** P<0.0001.

### The selective autophagy receptor p62 is recruited to staphylococci ingested by neutrophils

Selective autophagy has been shown to be involved in *S. aureus* infection of non-professional phagocytes [22], where evidence of bacterial ubiquitination and p62 recruitment was demonstrated. To confirm whether p62 is recruited to *S. aureus* in neutrophils, we used a *Tg(lyz:GFP-p62)i330* line (Gibson and Johnston, manuscript in preparation) hereafter called *lyz*:GFP-*p62*, in which the autophagy receptor protein p62 is fused to GFP under the neutrophil-specific *lyz* promoter. The *lyz*:GFP-*p62* larvae were infected with mCherry-expressing *S. aureus* and subjected to spinning disk confocal imaging. Within *S. aureus*-infected neutrophils, p62 commonly colocalises with intracellular bacteria (Fig. 7a) and also with apparent bacteria-containing vesicles (Fig. 7b). The p62-*S.aureus* colocalisation in neutrophils was observed in approximately 60% of infected neutrophils (Fig. 7c) although the pattern of p62 decoration differed from Lc3 (Fig. 2b). In contrast to Lc3 (Fig. 5), significantly less neutrophil p62-*S.aureus* associations were observed when heat-killed bacteria were used (Fig. 7c) suggesting that live bacteria are required to possibly damage and/or escape the phagosomes and recruit xenophagy receptors such as p62. Therefore, these results suggest that xenophagy might also occur within *S. aureus*-infected neutrophils and this effect is downstream of the initial recruitment of Lc3 to phagosomes.

**Figure 7:**
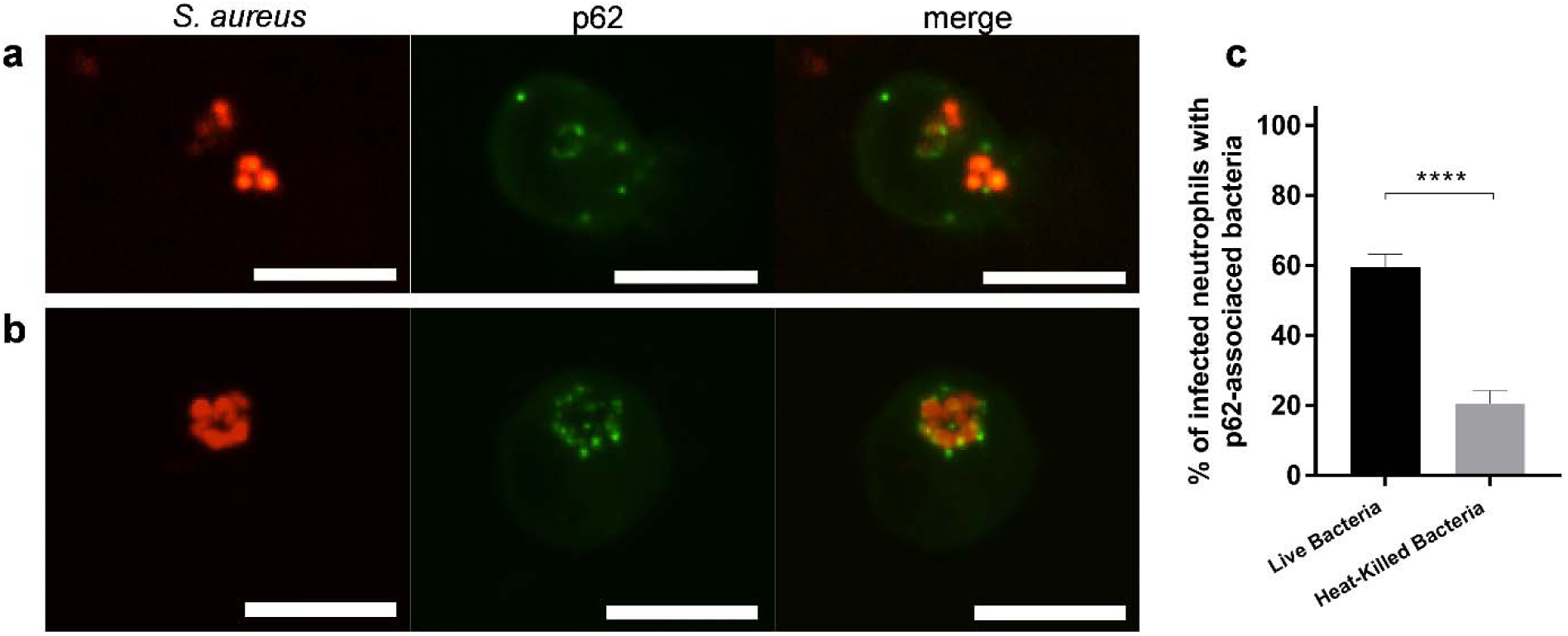
*S. aureus* within neutrophils is targeted by selective autophagy receptor p62. a, b. Confocal photomicrographs of p62-mediated response at 1 hpi in live *lyz*:p62-GFP embryos infected with mCherry-labelled *S. aureus*. The fusion p62-GFP protein colocalise with intracellular bacteria (a) or with apparent vesicles containing the bacteria (b). The images shown are representative of three independent experiments. Scale bars represent 10 µm.
c. Quantification of p62 associations with intracellular *S. aureus* at 2 hpi within neutrophils of *lyz*:p62-GFP infected with mCherry-labelled *S. aureus*. Data are shown as mean +/− standard error of the mean (SEM) obtained from three independent experiments. **** P<0.0001.

We also hypothesised that if Lc3-assocciated phagosomes (LAPosomes) formed within infected neutrophils are damaged by staphylococci, suppression of LAP would lead to reduction of p62 associations with intracellular bacteria. In order to inhibit LAP, *cyba* knockdown was performed and the formation of p62-positive structures associated with intracellular bacteria quantified. Indeed, we observed significantly less p62 association with intracellular bacteria in neutrophils, when LAP was blocked by *cyba* knockdown, suggesting that p62 association with bacteria is downstream of LAP (Fig. 8).

**Figure 8:**
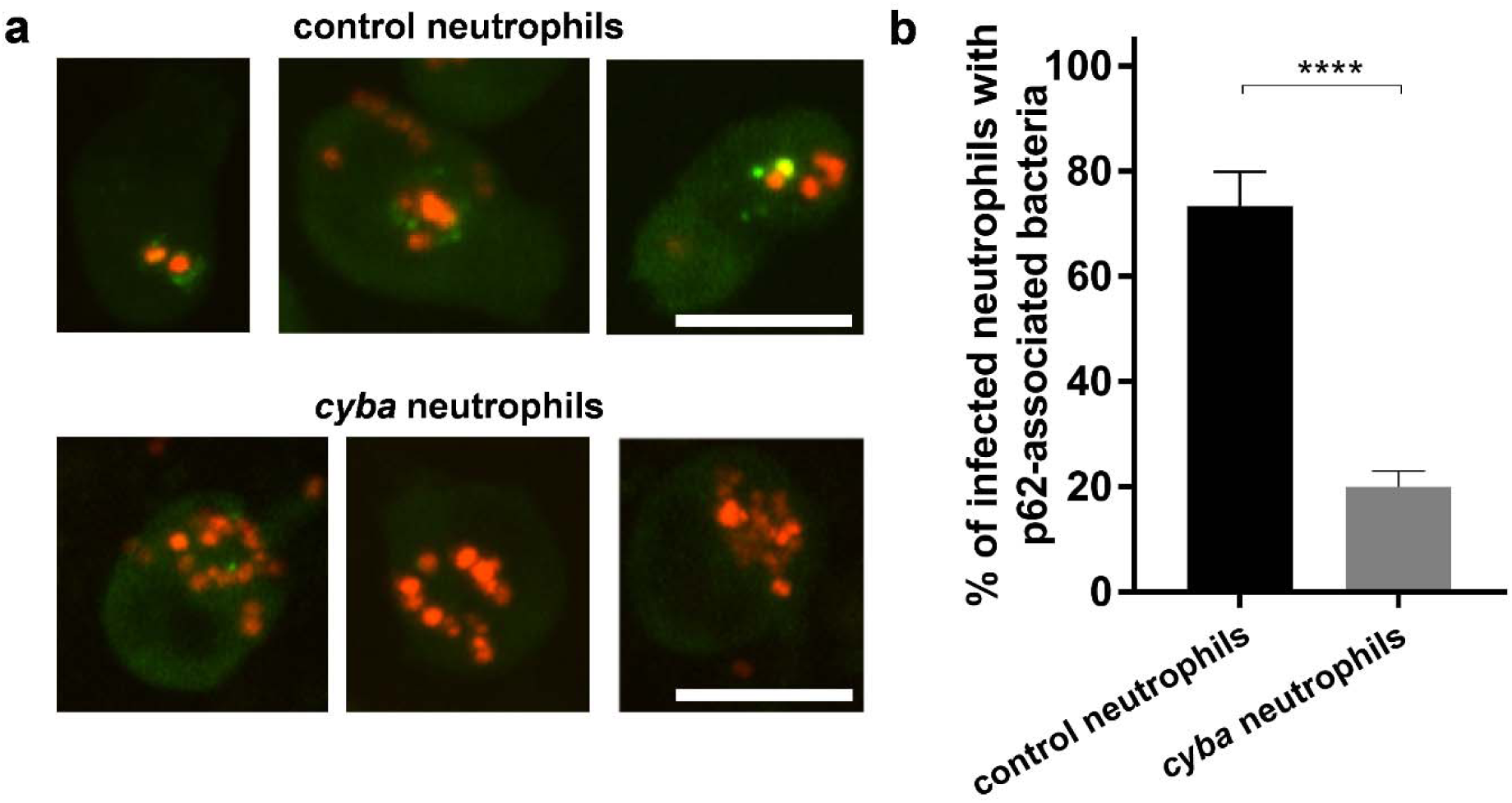
Diminished p62 recruitment to *S. aureus* in LAP-deficient neutrophils. a. Confocal photomicrographs of p62-mediated response at 2 hpi in live control (top panel) and *cyba* knockdown (bottom panel) *lyz*:p62-GFP embryos infected with mCherry-labelled *S. aureus*. The images shown are representative of three independent experiments. Scale bars represent approximately 10 µm.
b. Quantification of p62 associations with intracellular *S. aureus* at 2 hpi within neutrophils of *lyz*:p62-GFP infected with mCherry-labelled *S. aureus*. Data are shown as mean +/− standard error of the mean (SEM) obtained from three independent experiments. **** P<0.0001.

### Lc3 recruitment to *S. aureus* phagosomes is independent of the selective autophagy receptor p62

Neumann *et al.* have previously proposed that anti-staphylococcal p62-dependent xenophagy is protective to non-professional phagocytes infected by *S. aureus*, although to limited extent [22]. Similarly, our recent work in zebrafish has revealed that p62 is beneficial to zebrafish larvae infected by *S. aureus* (Gibson and Johnston, manuscript in preparation). To extend our understanding of the role of p62 in *S. aureus*-infected macrophages and neutrophils, we studied the effect of p62 deficiency on Lc3 recruitment to *S. aureus* in macrophages and neutrophils. The efficacy of knockdown using a p62 splice morpholino [40] was confirmed by RT-PCR (Fig. S9) and zebrafish embryos were subsequently infected with *S. aureus*. We found that the formation of Lc3-positive phagosomes in both macrophages and neutrophils was p62-independent as morpholino-mediated *p62* knockdown did not lead to reduction of Lc3-*S.aureus* association (Fig. 9a-c), further supporting that the initial Lc3-mediated response is indeed LAP and is p62-independent. However, in neutrophil enriched, macrophage-depleted larvae, loss of p62 caused mild but statistically significant increased susceptibility to *S. aureus* suggesting that p62 mediated processes, perhaps occurring following damage to the LAPosomes, are protective for *S. aureus*-infected neutrophils (Fig. 9d). Taken together, we propose that *S. aureus* exploits the autophagic response in neutrophils to establish an intracellular niche in LAPosomes, while p62-dependent selective autophagy in neutrophils may counteract the intracellular growth of the pathogen at later stages of infection.

**Figure 9.**
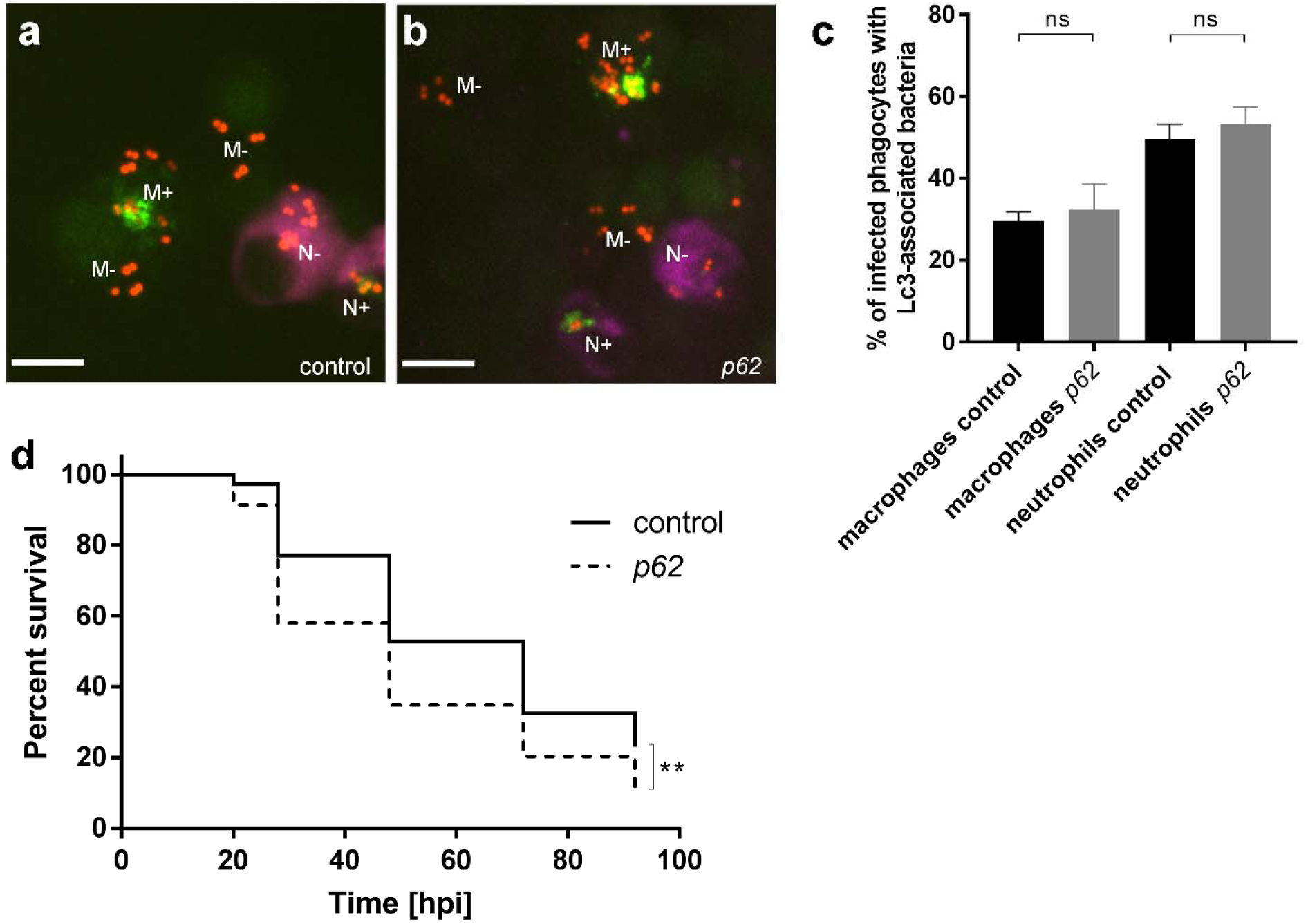
Loss of p62 leads to increased susceptibility to *S. aureus* infection. a, b. Confocal photomicrographs shown as maximum intensity projections of the Lc3-mediated response at 1 hpi within infected macrophages and neutrophils of control (a) and *p62* knockdown (b) CMV:GFP-Lc3 embryos infected with mCherry-labelled *S. aureus*. Embryos were fixed at 1 hpi and chemically stained for Mpx activity (TSA, magenta). TSA-negative macrophages are seen containing bacteria with (M+) or without (M-) Lc3 aggregates. TSA-positive neutrophils contain bacteria with (N+) and without Lc3 aggregates (N-). The images shown are representative of three independent experiments. Scale bars represent 10 µm.
c. Quantification of Lc3 associations with intracellular *S. aureus* at 1 hpi within infected macrophages and neutrophils of control and *p62* knockdown CMV:GFP-Lc3 embryos. Data are shown as mean +/− standard error of the mean (SEM) obtained from three independent experiments. ns – not significant.
d. Survival of *irf8* only or *irf8* + *p62* knockdown zebrafish larvae following intravenous injection with S. aureus at 30 hpf (n≥25). This result is representative of three independent experiments. ** *P*<0.01.

## DISCUSSION

*Staphylococcus aureus* has been shown to elicit an autophagic response in non-professional phagocytes as well as dendritic cells and macrophages [21,22,41,42], whilst autophagic responses to *S. aureus* in neutrophils have not been studied to date. To be able to utilise the autophagic machinery as a potential therapeutic target, several aspects of this host response need to be determined. First, what is the nature of the autophagic process targeting *S. aureus*, i.e. xenophagy or LAP? Secondly, what are the functional consequences of the autophagic response on different types of infected host cells? Do these processes promote bacterial pathogenesis or clearance within professional phagocytes infected with *S. aureus*? Using an established model of *S. aureus* infection in larval zebrafish, we demonstrate in this study that the autophagic machinery in neutrophils contributes to staphylococcal pathogenesis and that inhibition of this response improves host resistance.

Our findings relating to neutrophil function in the context of a whole organism are consistent with previous reports performed on *in vitro* cultured non-professional phagocytes [21,23], wherein the autophagic machinery of host cells provided a niche for staphylococcal dissemination. We propose that, at early stages of *S. aureus* infection, infected phagocytes undergo Lc3-associated phagocytosis. This is because this Lc3-mediated response occurs rapidly post-infection (within 1 hpi), the Lc3 signal labels the membrane of spacious phagosomal compartments, Lc3 recruitment does not require live staphylococci, and Lc3 recruitment is independent of p62, suggesting damage to the phagosomal membrane is not required. Moreover, the formation of *S. aureus*-containing Lc3-positive phagosomes required ROS production by phagocyte NADPH oxidase, another hallmark of LAP [18,19,43].

In addition to demonstrating the NADPH oxidase-dependent recruitment of Lc3 in *S. aureus* infected neutrophils, we observed that the Lc3 association with *S. aureus*-containing phagosomes in neutrophils was prolonged up to at least 6 hpi in comparison to the response observed in macrophages. Similar results were observed by Huang *et al.* where neutrophils treated with IgG-coated beads also showed LC3 associations with phagosomes in a DPI-sensitive manner for extended periods of time [43]. Therefore, it seems that the recognition and subsequent clearance of LC3-labelled phagosomes through the autophagic pathway can be strongly inhibited in neutrophils, which can be utilised by intracellular microbes for pathogenesis. Using a novel zebrafish transgenic line *lyz*:RFP-GFP-Lc3 to study the Lc3 association specifically in neutrophils, we observed formation of spacious Lc3-positive vesicles. These are similar to previously observed spacious Listeria-containing phagosomes (SLAPs) of mouse macrophages, the LC3-positive structures associated with persistent disease [19,44]. Our live imaging studies in zebrafish are consistent with electron microscopy data of murine neutrophils, where virulent staphylococci, but not an attenuated *sar* mutant strain, also induced the formation of spacious phagosomes [8]. In another study performed on murine bone marrow-derived dendritic cells, it was demonstrated that *S. aureus* is able to inhibit autophagic flux and chemical inhibition of the autophagic response by 3-MA reduced cytotoxicity caused by phagocytosed staphylococci [45], hence resembling the response observed in neutrophils in our study.

Importantly, we show a pronounced difference in Lc3-mediated response between macrophages and neutrophils, where more neutrophils with staphylococci-containing LAPosomes were observed than macrophages and this difference was especially more apparent at later stages of infection. Therefore, it is likely that LAP also occurs in zebrafish macrophages, but this response might be quickly resolved leading to subsequent loss of Lc3 association with the phagosome upon fusion with lysosomes. Indeed, using a *Salmonella* infection model, we have recently shown that zebrafish macrophages are able to mount a LAP response that promotes bacterial clearance [33]. Similar to our results, a study using murine RAW264.7 macrophages infected by staphylococci showed that the Lc3-mediated response peaks at 1 hpi and subsequently drops until 4 hpi [42]. Another recent work on RAW264.7 macrophages revealed that only a fraction of phagosomes containing *S. aureus* were Lc3-positive within 12 hours of infection, indicating a low level of autophagic response and little phagosomal damage caused by staphylococci within infected macrophages [46]. Thus, it appears that LAP might also occur in infected macrophages, but unlike in neutrophils, this response is rapidly processed by the autophagic flux and hence a time-dependent loss of Lc3-bacteria associations is generally observed.

Chronic Granulomatous Disease (CGD) patients are more susceptible to *S. aureus* infection, predominantly manifested by liver abscesses and skin and soft tissue infections [47] but not septicaemia. The exact reason why CDG patients are more prone to these staphylococcal infections is not fully understood. Although neutrophils are considered major ROS-producing phagocytes, it has been recently shown that the effect of NADPH oxidase inhibition is more pronounced in macrophages as *Ncf1* (*p47phox*) mutant mice with ectopic expression of *Ncf1* in monocytes/macrophages are protected from experimental *S. aureus* systemic infection [48]. Therefore, the exact role of NADPH oxidase in neutrophils needs to be evaluated in the context of both local and systemic staphylococcal infection. In addition, our results are in line with a recent study which has shown that human neutrophils devoid of phagosomal ROS production due to NADPH oxidase mutation, although not being able to kill intracellular bacteria, are fully capable of containing staphylococci in a 3D matrix *in vitro* model. Moreover, a subset of human neutrophils with highest acidification rate has been found to be the most efficient in containing the staphylococcal infection [49]. Perhaps this enhanced ability to contain staphylococci is host-protective in our bacteraemia model of infection and the host-detrimental effect of LAP in *S. aureus*-infected zebrafish is likely due to the formation of a non-acidified niche driven by live *S. aureus*, which could subsequently lead to neutrophil lysis and bacterial escape.

In addition to LAP, we observed that p62 also targets *S. aureus* within infected zebrafish neutrophils and loss of p62 led to mildly increased susceptibility, suggesting that xenophagy might be also involved and plays a protective role in *S. aureus* infection of neutrophils. Similarly, an *in vitro* work by Neumann *et al.* reported that *S. aureus* is decorated by p62 and encapsulated within autophagosomes of non-professional phagocytes, and that blocking autophagosome formation by ATG5 knockout leads to higher bacterial loads [22]. On the other hand, we have observed that blocking LAP by *cyba* knockdown leads to reduced p62 recruitment to bacteria in neutrophils, suggesting that formed LAPosomes subsequently undergo membrane damage induced by live staphylococci and therefore injection with heat-killed bacteria also led to diminished p62 recruitment and possible xenophagy. Mitchel *et al.* have observed similar phenomenon in macrophages infected by *Listeria monocytogenes* where bacteria were initially encapsulated within LC3-positive phagosomes and subsequent damage of the vacuoles triggers xenophagy [50]. In conclusion, our results suggest a dual role for the autophagy machinery in neutrophils, with the formation of LAPosomes facilitating the intracellular life stage of *S. aureus* and xenophagy providing partial protection.

Therefore, we propose the following model of the fate of staphylococci within neutrophils (Fig. 10). Bacteria are internalised by neutrophils and trapped within LC3-associated phagosomes which are triggered by phagosomal NADPH oxidase. These LAPosomes do not get acidified, allowing internalised *S. aureus* to damage the phagosomal membrane. Bacteria or damaged phagosomal membrane may then be detected by the selective autophagy receptor protein p62 which might lead to formation of autophagosomes containing staphylococci and pathogen inactivation. The inability to sequester all bacteria from damaged LAPosomes or inhibition of autophagy flux could lead to subsequent bacterial dissemination. Taken together, the observed antagonistic role of the autophagic machinery (LAP vs. p62-mediated response) within *S. aureus*-infected neutrophils may explain the conflicting reports on anti-staphylococcal autophagy. Clarification of the molecular mechanisms how *S. aureus* is engaged in LAP awaits future study, which may provide new insights for therapeutic strategies to fight this intracellular pathogen.

**Figure 10.**
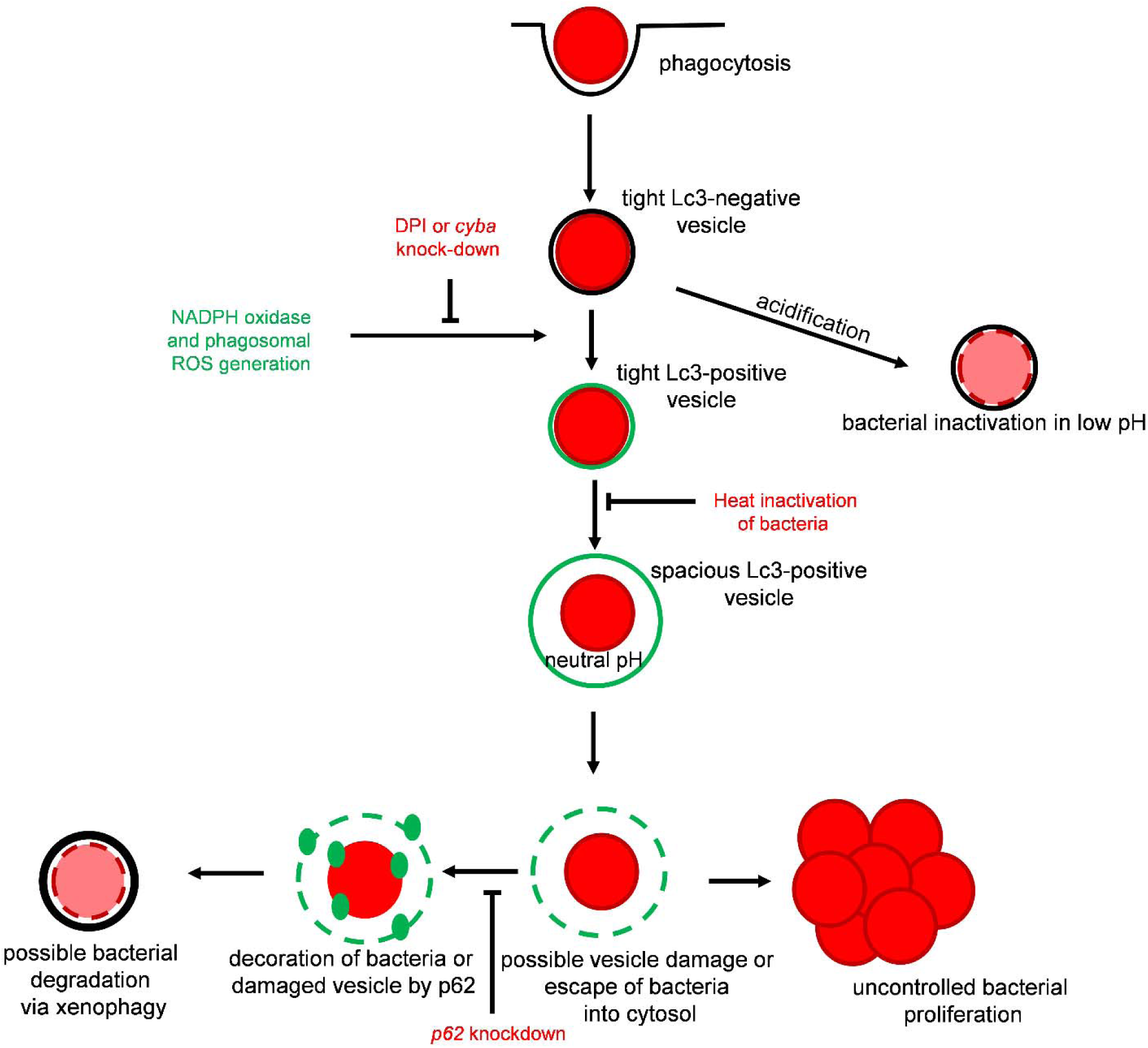
Graphical Abstract: Proposed model of fate of *S. aureus* within neutrophils. Staphylococci are internalised by neutrophils and trapped within LC3-associated phagosomes (LAPosomes) triggered by NADPH oxidase. LAPosomes do not get acidified and provide a replication niche for internalised *S. aureus*, which eventually will damage the phagosomal membrane and escape into the cytoplasm. This is sensed by selective autophagy machinery which leads to formation of autophagosomes containing staphylococci and leading to pathogen inactivation.

## MATERIALS AND METHODS

### Zebrafish lines and maintenance

Zebrafish adults and embryos were handled in compliance with local animal welfare regulations and maintained according to standard protocols (zfin.org) in compliance with international guidelines specified by the EU Animal Protective Directive 2010/63/EU. Existing lines were London wild-type (LWT), *Tg(CMV:EGFP-map1lc3b)zf155* [27] and *Tg(lyz:GFP-p62)i330* (Gibson and Johnston, manuscript in preparation). For the generation of the *Tg(lyz:RFP-GFP-Lc3)sh383* line, zebrafish RFP-GFP-Lc3 middle entry clone [34] was used to generate pDEST(*lyz*:RFP-GFP-Lc3). Embryos of wild type zebrafish lines (LWT) were injected with transgenic expression construct together with the tol2 transposase. Positive embryos were selected for mosaic expression under a Leica fluorescence dissecting microscope by GFP heart marker. Selected embryos were raised to adulthood and after 3 month sexually mature fish were screened by outcrossing to LWT fish. The offspring of potential founders were screened for transgene expression. Embryos were incubated in E3 medium at 28.5°C according to standard protocols [51].

### Bacterial Cultures and Infection experiments

*Staphylococcus aureus* SH1000 expressing mCherry [28] was cultured in brain heart infusion (BHI) broth medium (Sigma) at 37°C supplemented with tetracycline at 5 µg/ml. Zebrafish larvae at 30 hpf (hours post fertilisation) were microinjected into the circulation with bacteria as previously described [25]. Briefly, anaesthetized larvae were embedded in 3% w/v methylcellulose and injected individually using microcapillary pipettes filled with the bacterial suspension of known concentration. For macrophage depletion, clodronate liposomes were injected at 26 hpf as previously described [38]. Following infection, larvae were observed frequently up to 120 hpf, and numbers of dead embryos recorded at each time point.

### Determination of *in vivo* bacterial load

At various times post infection, living zebrafish larvae were anesthetized and individually transferred with 100 µl of E3 medium into 0.5 ml tubes containing 1.4 mm ceramic beads and homogenized using a Precellys 24-Dual homogenizer (Peqlab). The homogenates were serially diluted and plated on BHI agar to determine *S. aureus* CFU numbers. Bacterial load was also determined for dead larvae at each time point.

### Morpholino knockdown and RT-PCR for morpholino efficacy verification

Morpholino oligonucleotides (Gene Tools) were dissolved in MilliQ water to obtain the required concentrations. 1 nl volume of morpholino was injected into the yolk of 1-4 cell stage zebrafish embryos using a microinjector. Standard control morpholino (Gene Tools) was used as a negative control. The *irf8* [30], *cyba* [37] and *p62* [40] morpholinos were used at the previously published concentrations. The *p62* knockdown was verified by RT-PCR with a pair of primers flanking the splicing event between the first intron and the second exon (i1e2) as described before [40].

### TSA staining

Infected embryos were fixed in ice-cold 4% (w/v) paraformaldehyde (PFA) in PBS-TX (PBS supplemented with 0.5 % of Triton X-100) overnight at 4°C. Fixed embryos were washed in PBS-TX twice. Peroxidase activity was detected by incubation in 1:50 Cy5-TSA : amplification reagent (PerkinElmer, Waltham, MA) in the dark for 10 min at 28°C followed by extensive washing in PBS-TX.

### Staining of *S. aureus* with pH-sensitive dyes

The pHrodo Red and Fluorescein-5-EX S-ester dyes (Life Technologies) were dissolved in DMSO to the final concentrations of 2.5 mM and 16.95 mM, respectively. 0.5 μl of pHrodo Red and 1.5 μl Fluorescein was added to 200 μl of bacterial suspension in PBS pH 9 and then mixed thoroughly. The mixture was incubated 30 min at 37 °C with gentle rotating. To remove the excess of the dyes, bacteria were washed during 3 step procedure: addition of 1 ml of PBS pH 8, 1 ml of Tris pH 8.5, again 1ml of PBS pH 8; followed by 2 min of centrifugation in 12000 g and gentle removal of the supernatant. After washing, bacterial pellet was resuspended in 200 μl of PBS pH 7.4 and proceeded to microinjections of zebrafish embryos.

### Imaging and Image analysis

Live anesthetized or PFA-fixed larvae were mounted in 1% (w/v) low-melting-point agarose solution in E3 medium. For live larvae, images were acquired using the UltraVIEW VoX spinning disk confocal microscope (Perkin Elmer) with Olympus 40x UPLFLN oil immersion objective (NA 1.3). For fixed samples, images were acquired using Leica TCS SPE laser scanning confocal microscope with 63x HC PL APO water immersion objective (NA 1.2).

For quantification of the autophagic response within infected phagocytes, for each embryo, a total number of observable infected phagocytes were manually determined through the z-stacks of acquired images. Among these total observable infected phagocytes number of infected phagocytes with GFP-Lc3 signal were enumerated, and percentage of Lc3-positive phagocytes over total observable phagocytes was determined for each embryo. Maximum projections were used for representative images. No non-linear normalizations were performed.

### Statistical Analysis

Survival experiments were evaluated using the Kaplan-Meier method. Comparisons between curves were made using the Log Rank (Mantel Cox) test. Quantifications of percent Lc3-positive phagocytes or CFU counts were determined for significance with unpaired parametric t-test for 2 groups and with one-way ANOVA for multiple groups, corrected for multiple comparisons. Analysis was performed using Prism version 7.0 (GraphPad). Statistical significance was assumed at P values below 0.05.

## ACKNOWLEDGMENTS

We thank Dan Klionsky (University of Michigan) for the CMV:GFP-Lc3 zebrafish line. We are grateful to all members of the fish facility teams at Institute of Biology Leiden and Bateson Centre for zebrafish care. T.K.P. was supported by an individual Marie Curie fellowship (PIEF-GA-2013-625975) and by AMR cross-council funding from the MRC to the SHIELD consortium “Optimising Innate Host Defence to Combat Antimicrobial Resistance” MRNO2995X/1. J.J.S. was a Marie Curie fellow in the Initial Training Network FishForPharma (PITN-GA-2011-289209), both funded by the 7th Framework Programme of the European Commission. S.M. was supported by a fellowship from the Higher Education Commission of Pakistan and the Bahaudin Zakriya University, Multan. Live imaging used the Wolfson Light Microscopy Facility (supported by MRC grant MR/K015753/1).

---

Figure S1. Different kinetics of the Lc3-mediated response in macrophages and neutrophils infected by *S. aureus*

a. Confocal photomicrographs shown as maximum intensity projections of control (top panel) and *irf8* knockdown (bottom panel) CMV:GFP-Lc3 embryos infected with mCherry-labelled *S. aureus*. Embryos were fixed at 6 hpi and chemically stained for Mpx activity (TSA, magenta). The images shown are representative of three independent experiments. Scale bars represent 10 µm.
b. Quantification of Lc3 associations with intracellular *S. aureus* at 6 hpi within neutrophils. Data are shown as mean +/− standard error of the mean (SEM) obtained from three independent experiments. ns – not significant.

Figure S4: Formation of NADPH oxidase-dependent Lc3-positive vesicles containing *S. aureus* in neutrophils is detrimental for the infected host

a. Survival of control (DMSO) and DPI-treated *irf8* knockdown zebrafish larvae following intravenous injection with *S. aureus* at 30 hpf (n≥25). This result is representative of three independent experiments. * *P*<0.05.
b. Survival of macrophage-depleted (clodronate-treated) control and *cyba* knockdown zebrafish larvae following intravenous injection with *S. aureus* at 30 hpf (n≥25). This result is representative of three independent experiments. *** *P*<0.001.
c. Survival of control and *cyba* knockdown zebrafish larvae following intravenous injection with *S. aureus* at 30 hpi (n≥25). This result is representative of three independent experiments. ns – not significant.

Figure S9. Loss of p62 leads to increased susceptibility to *S. aureus* infection

Electrophoresis gel scan of reverse transcription polymerase chain reaction (RT-PCR) products of control and p62 knockdown zebrafish at 2 dpf used to determine the efficacy of *p62* splice morpholino.

