## Supplementary figures and images for "The autophagic response to *Staphylococcus aureus* provides an intracellular niche in neutrophils"

### Figure S1

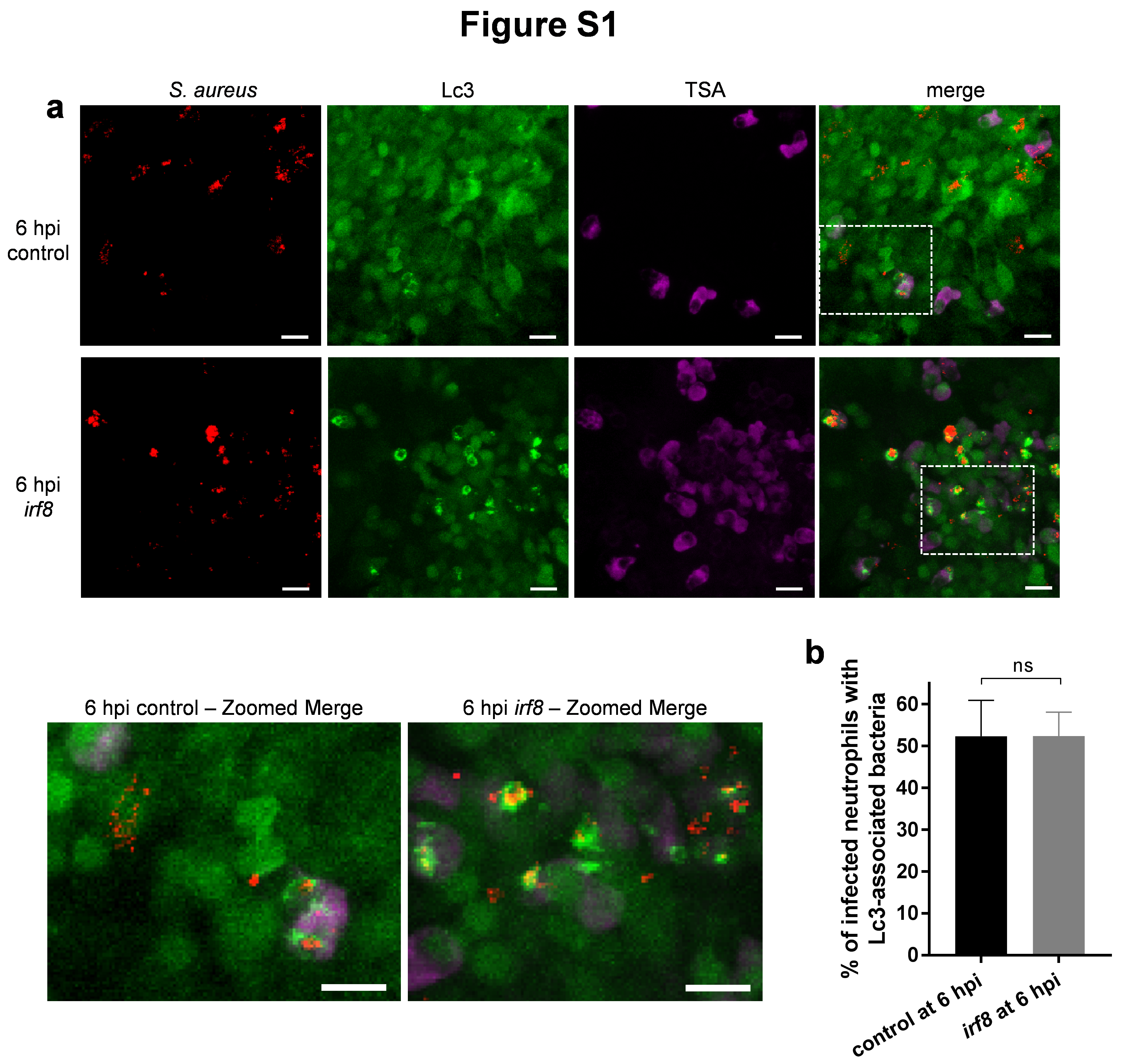

### Figure S4

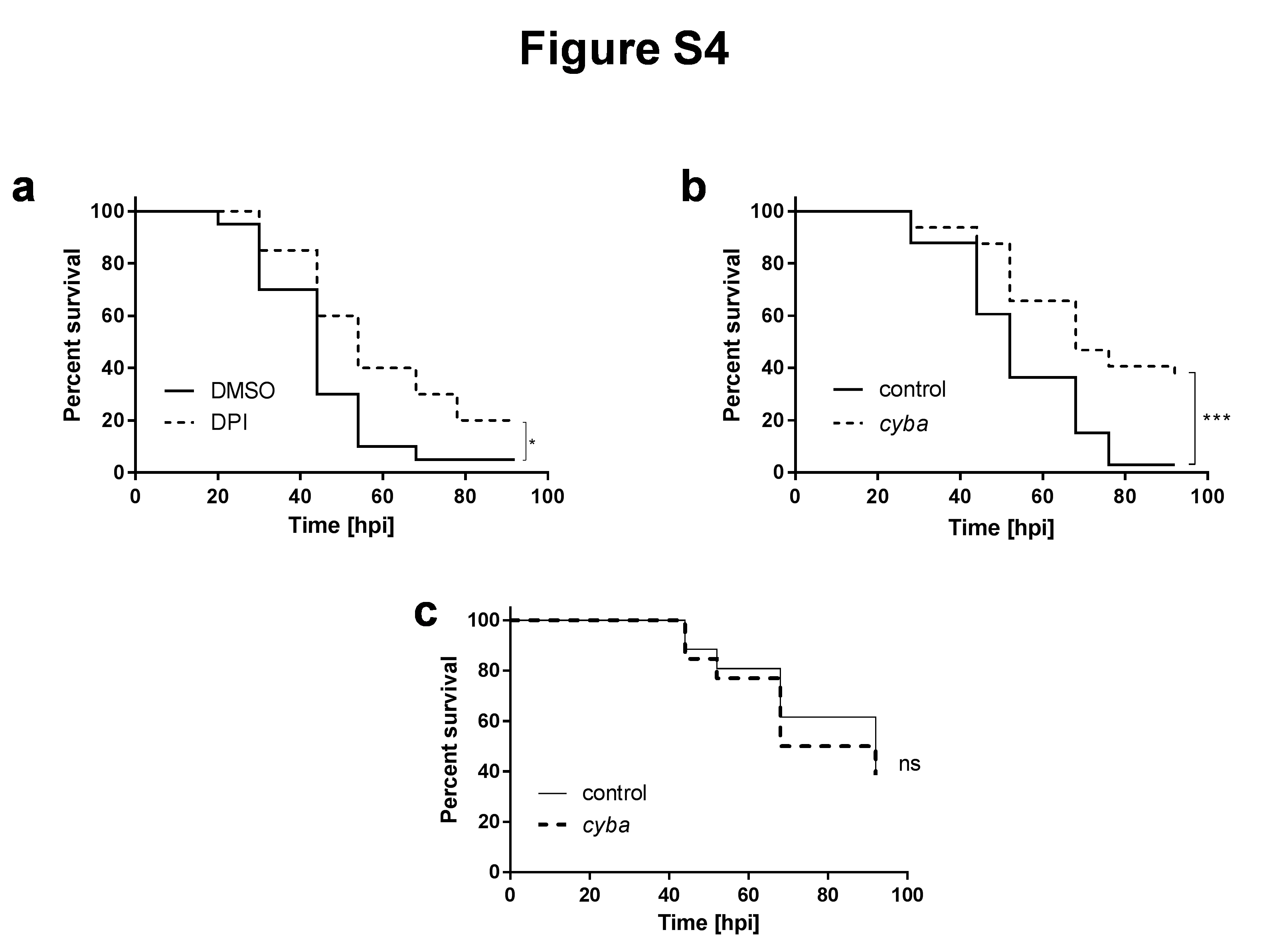

### Figure S9

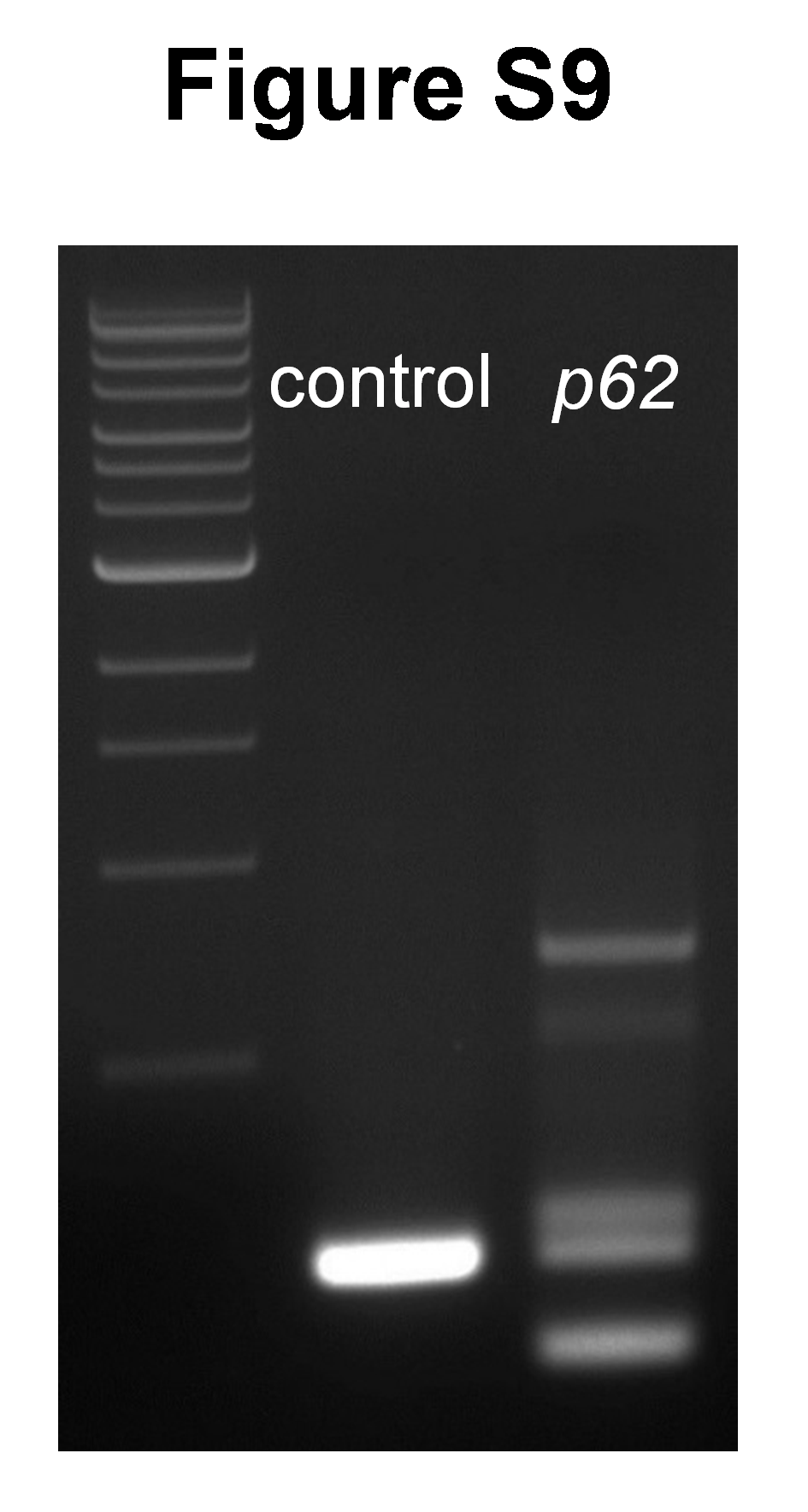
